# Early Adversity Selectively Reshapes the Somato-Cognitive Action Network in the Developing Brain

**DOI:** 10.64898/2026.07.17.739165

**Authors:** Chase Antonacci, Eugenia Giampetruzzi, Gracie Grimsrud, Sanju Koirala, Jessica P. Uy, Jocelyn A. Ricard, Yoonji Lee, André Zugman, Booil Jo, Kilian M. Pohl, Russell A. Poldrack, Ian H. Gotlib

## Abstract

Early adversity is among the most potent environmental influences on the developing brain; yet whether it remodels the areal layout of cortical networks, and along which dimensions, is unknown. In a longitudinal sample of 4,525 youth aged 9-18, we derived individualized cortical parcellations and related the topography of 15 networks to 10 data-driven dimensions of adversity. Greater exposure to threat, but not deprivation or unpredictability, predicted dose-dependent expansion of the somato-cognitive action network (SCAN), an effect nearly seven times larger than for any other network. The SCAN expanded toward the sensorimotor pole, encroached on neighbors, and shifted functional connectivity toward sensorimotor and away from control systems. Expansion mediated the association between threat and poorer cognition, replicating across waves, within children, and out-of-sample. Rather than diffusing across many systems, the imprint of adversity on the developing cortex converges on the territory of a single integrative network that couples brain to body.

## Introduction

Early life adversity (ELA) is among the most potent environmental influences on the developing brain, with lasting consequences for cognition, emotional functioning, and mental health across the lifespan^1–3^. Adverse experiences such as violence, maltreatment, poverty, and neglect have been associated with poorer cognitive and academic outcomes^4,5^ and account for up to a third of adult psychiatric disorders worldwide^6,7^. Over the past decade, researchers have linked adversity both to alterations in brain structure and to aberrant functional connectivity within and between large-scale cortical networks^8–11^. Far less is known, however, about how adversity shapes the functional topography of the cortex itself – that is, the spatial layout of its networks and how much area each cortical network occupies. Critically, emerging research suggests that cortical topography varies across individuals and is related to differences in cognitive ability^12,13^ and risk for mental health difficulties^14,15^. Whether the environment in which a child is raised influences this organization, and which dimensions of adversity exposure matter most, have been largely neglected in the study of how early experience becomes biologically embedded.

Leveraging advances in precision functional mapping of the brain, researchers have established that the spatial organization of these networks (i.e., their functional topography) is idiographic and stable over time^12,16,17^. Moreover, interindividual variation in topography has been found to be heritable^18^ and to predict cognition and behavior more accurately than do spatial maps derived from group averages^19,20^. The boundaries of these networks are refined across adolescence, most extensively in the association cortex^19,21,22^, and can shift as a function of mental illness, illustrated by expansion of the salience network in adults with depression^14^ and with obsessive compulsive disorder^23^. The recent discovery of the somato-cognitive action network (SCAN), a system interleaved within the motor cortex that integrates goal-directed action with autonomic and physiological control^24^, indicates that our understanding of functional organization is still being refined. Together, these findings establish functional topography as a stable but individually variable phenotype, one in which the cortex apportions territory among networks in ways that change with age and bear on cognition and mental health.

Given that adolescence is a period of pronounced cortical plasticity, a central question is whether the early environment leaves an imprint on this organization. Dimensional formulations of early adversity hold that the type of exposure a child faces has consequences for brain and behavior. Specifically, experiences of threat (experiences involving harm or its anticipation, such as violence and abuse) and deprivation (the absence of expected cognitive and social inputs, such as neglect and poverty) have been posited to engage partly separable neural systems^25–28^, with environmental unpredictability posited to be a third dimension of adversity^29^. We do not yet know, however, whether these dimensions map onto distinct features of cortical topography. In this context, researchers have begun to link aspects of the early environment to individualized network organization; in fact, a general environmental factor that aggregates children’s exposures into a single composite has been found to predict both network topography and overall cognition in youth^30^. Aggregating exposures this way, however, masks which domains of adversity might account for the effect or which networks are reorganized. In a more targeted study, Butler et al.^31^ examined whether exposure to violence expands the salience network in adolescence, an intuitive candidate network given its role in threat processing, but found no significant association. Therefore, characterizing whether and how the environment shapes cortical topography requires an approach that resolves distinct dimensions of adversity, surveys the entire cortex, and traces how this organization unfolds across adolescence.

The present study was designed to test whether early adversity alters the spatial organization of functional brain networks. We derived individualized functional network parcellations in a large sample of youth from the Adolescent Brain Cognitive Development (ABCD) Study who were followed across four waves from ages 9-18 years. Given that almost no prior work has examined how the environment shapes network topography, we did not hypothesize effects of a specific dimension of adversity on a particular network. Instead, we conducted a whole-cortex screen, relating the cortical territory of all 15 networks in each child to 10 data-driven dimensions of adversity that together summarize the full range of environmental exposures assessed in ABCD, which we then grouped into threat, deprivation, and unpredictability composites reflecting the classic taxonomy. Findings converged on a single, unanticipated result: threat, but not deprivation or unpredictability, predicted pronounced expansion of the SCAN, an effect nearly seven times larger than that for any other network and graded by exposure. This expansion concentrated toward the sensorimotor pole of the cortical hierarchy, encroached on adjacent networks, and was accompanied by a shift in functional coupling toward sensorimotor and away from control systems. Importantly, it mediated the relation between threat and crystallized cognition, an effect that replicated across waves, persisted within children over time, and predicted cognition in held-out youth. These findings demonstrate, for the first time, that a specific dimension of early adversity predominantly reconfigures a single integrative brain network, identifying a distinct mode of biological embedding in the developing cortex.

## Results

### Participant Characteristics

The primary analytic sample was composed of 4,525 children (4,058 families, 21 sites) with usable baseline resting-state fMRI, individualized network parcellations, and complete ELA data; full sample characteristics are reported in Table 1. Seated within the supplementary and cingulate motor, insular, and opercular cortex (Fig. 1b), the SCAN occupied a median of 2.5% of the cortical surface (IQR 1.9-3.3%; Fig. 1d), making it the second smallest of the 15 networks.

**Table 1.** Sample Characteristics.

| **Characteristic** | ***N (%) or M (SD)*** |
| --- | --- |
| Total participants | 4,525 |
| Unique families | 4,058 |
| Participants with a sibling in sample | 923 (20.4%) |
| Study sites | 21 |
| Age at baseline (years) | 9.98 (0.63) |
| **Sex** |  |
| Male | 2,171 (48.0%) |
| Female | 2,354 (52.0%) |
| **Race/ethnicity** |  |
| White | 2,627 (58.1%) |
| Hispanic | 700 (15.5%) |
| Other/Multiracial | 456 (10.1%) |
| Black/African American | 394 (8.7%) |
| Asian | 64 (1.4%) |
| Missing | 284 (6.3%) |
| **Household income** |  |
| ≥ \$100,000 | 2,327 (51.4%) |
| \$50,000–99,999 | 1,116 (24.7%) |
| < \$50,000 | 798 (17.6%) |
| Missing | 284 (6.3%) |
| **Parental education** |  |

| **Characteristic** | **N (%) or M (SD)** |
| --- | --- |
| Bachelor's degree | 1,384 (30.6%) |
| Postgraduate degree | 1,263 (27.9%) |
| Some college / associate | 1,115 (24.6%) |
| High-school diploma / GED | 336 (7.4%) |
| < High-school diploma | 143 (3.2%) |
| Missing | 284 (6.3%) |
| **Early adversity exposures**□ |  |
| Threat composite (z) | 0.00 (1.00) |
| Deprivation composite (z) | 0.00 (1.00) |
| Unpredictability composite (z) | 0.00 (1.00) |
| High-threat subgroup ($\geq +1$ SD) | 579 |
| Low-threat subgroup ($\leq -1$ SD) | 554 |
| **Imaging and motion QC (baseline rest)** |  |
| Mean framewise displacement (mm) | 0.118 (0.061) |
| Retained volumes post-scrubbing | 1,189 (184) |
| Retained rest data (minutes) | 15.9 (2.4) |
| **Scanner manufacturer** |  |
| Siemens | 3,012 (66.6%) |
| GE | 1,102 (24.4%) |
| Philips | 411 (9.1%) |
| **SCAN territory (% of cortex)** |  |
| Mean (SD) | 2.73 (1.12) |
| Median [IQR] | 2.51 [1.91-3.31] |
| **Cognitive and behavioral outcomes** |  |
| NIH-TB Crystallized, year 6 (n = 2,997) | 106.64 (15.83) |
| NIH-TB Fluid, year 6 (n = 2,934) | 113.91 (19.34) |
| CBCL Total Problems, year 6 (n = 2,665) | 13.47 (15.44) |
| CBCL Attention Problems, year 6 (n = 2,665) | 2.17 (3.01) |

| *Characteristic* | *N (%) or M (SD)* |
| --- | --- |
| CBCL DSM-5 ADHD, year 6 (n = 2,665) | 1.69 (2.32) |
| CBCL Thought Problems, year 6 (n = 2,665) | 1.13 (1.83) |

**Note.** Data are from the Adolescent Brain Cognitive Development (ABCD) Study (N = 4,525 children with usable baseline resting-state fMRI data and complete early life adversity factor scores). For categorical variables, values are n (%); for continuous variables, values are *M* (*SD*), except SCAN territory, for which the second row reports the median and interquartile range [IQR]. Baseline age ranged from 8.9 to 11.0 years. Race/ethnicity, household income, and parental education were missing for the same 284 participants (6.3%); percentages may not sum to 100 because of rounding. CBCL scores are raw problem-scale sums, and NIH-TB scores are age-corrected standard scores. CBCL = Child Behavior Checklist; NIH-TB = NIH Toolbox; SCAN = somato-cognitive action network; QC = quality control; FD = framewise displacement; GED = General Educational Development; IQR = interquartile range; *M* = mean; *SD* = standard deviation.
<sup>a</sup> Threat, deprivation, and unpredictability composites are standardized (z-scored) across the analytic sample; *M* (*SD*) = 0.00 (1.00) by construction, so the informative quantities are the $\geq +1$ SD and $\leq -1$ SD subgroup sizes. Higher scores indicate greater exposure.

**Figure 1:**
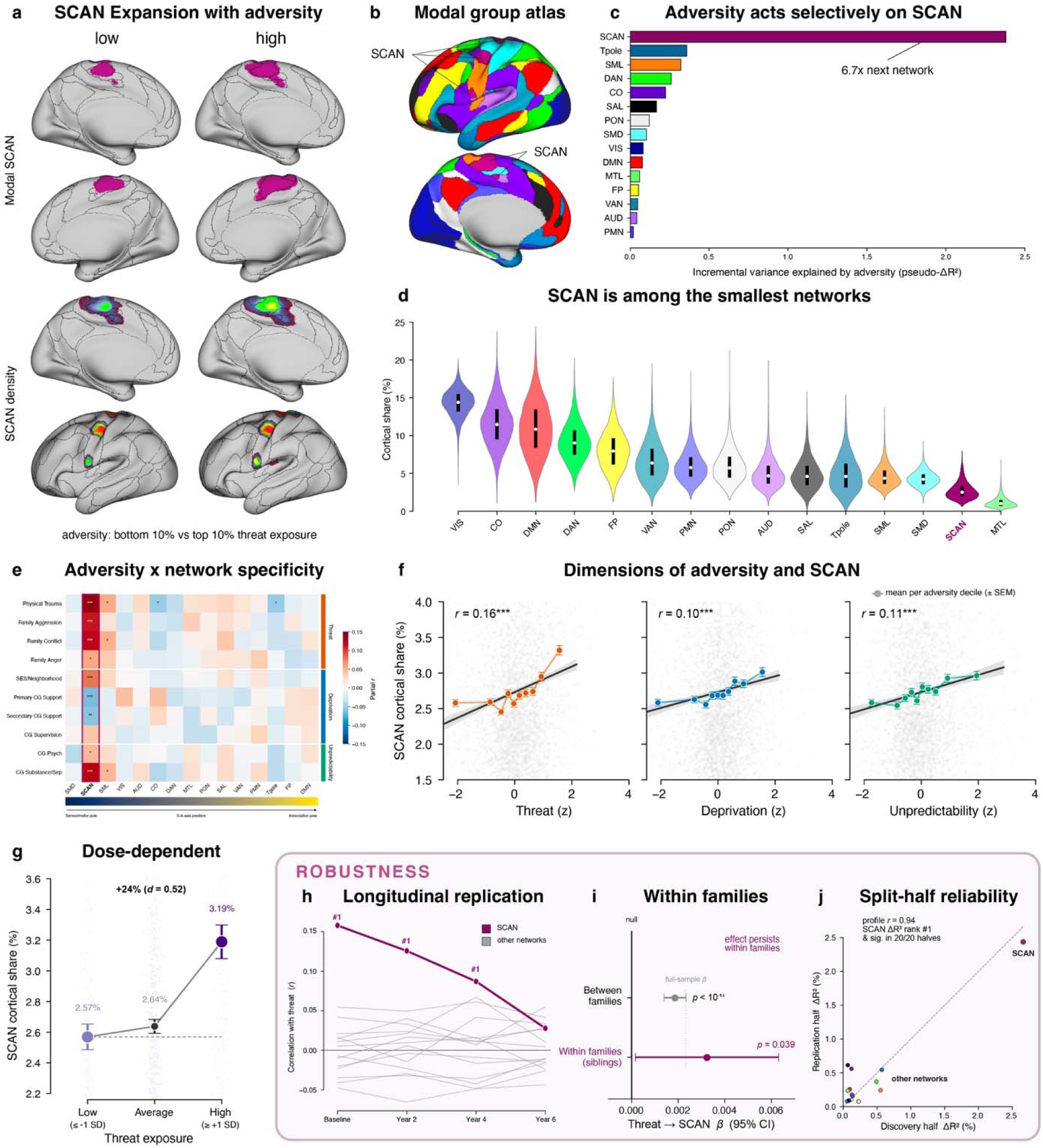
Early adversity selectively expands the SCAN network. **(a)** Group SCAN topography in youth with low (≤10th percentile) versus high (≥90th percentile) threat exposure, shown as the modal (winner-take-all) SCAN territory (top) and the continuous across-subject SCAN density (bottom), on medial and lateral surfaces. SCAN territory is visibly larger in high-threat youth. **(b)** Population modal atlas of all 15 cortical networks, for anatomical context. **(c)** Variance in network size uniquely explained by adversity (pseudo-ΔR²) across all 15 networks: the effect is concentrated on SCAN, 6.7x the next-ranked network. **(d)** Networks ranked by cortical share; SCAN is among the smallest networks (rank 14/15), making its outsized adversity sensitivity notable. **(e)** Partial correlations between 10 adversity subscales (grouped into threat, deprivation, and unpredictability) and all 15 networks; the SCAN column (boxed) shows the strongest and most consistent associations (* FDR *q* < .05; ** *q* < .01, *** *q* < .001). Networks are ordered along the sensorimotor-to-association (S-A) axis, placing SCAN near the sensorimotor pole. **(f)** SCAN cortical share increases with each adversity dimension – threat (*r* = 0.16), deprivation (*r* = 0.10), and unpredictability (*r* = 0.11), all *p* < .001; points are per-decile means (± SEM). **(g)** Dose-response: SCAN share rises from 2.57% in low-threat to 3.19% in high-threat youth, a 24% increase (*d* = 0.52). **(h)** Longitudinal replication: threat → network correlations across waves; SCAN (magenta) is the top-ranked network at baseline through Year 4 and attenuates by Year 6. **(i)** The threat → SCAN association persists within families, in a sibling/within-family model (β > 0, *p* = 0.039), not just between families (*p* = 1.09 × 10^-14^), controverting a purely between-family confound. **(j)** Split-half reliability: per-network adversity ΔR^2^ in independent discovery and replication halves; SCAN ranks #1 in both (profile *r* = 0.94; SCAN top-ranked in 20/20 splits). SCAN is shown in magenta throughout.

Baseline proportional SCAN size was comparable in female and male children (2.70% vs 2.77%; *d*=0.06, *p*=.054). Relative to the remainder of the ABCD baseline cohort, the analytic sample was modestly more likely to be female, White, and from higher-income and higher-education households (all bias-corrected^32^ Cramér’s *V*≤0.17), a selection pattern consistent with the well-documented association between MRI scan quality and sociodemographic factors^33^.

### Early Adversity Selectively Expands the Somato-Cognitive Action Network

We first tested whether early adversity is associated with an altered cortical functional network map, and with what anatomical specificity. Testing each of the 10 data-driven ELA factors against the cortical proportion of all 15 functional networks, we found that the associations converged on a single network. Nine of the ten dimensions were independently associated with the proportion of cortex allocated to the SCAN at baseline (all FDR *q*< 0.05), a consistency that was not observed for any other network (Fig. 1e). The strongest associations were with physical trauma (partial *r*=0.131, *q*<.0001, BF_10_=5.3×10^16^), caregiver substance use or separation (*r*=0.113, *q*<.0001, BF_10_=4.6×10^11^), family conflict (*r*=0.110, *q*<.0001, BF_10_=5.7×10^10^), and family aggression (*r*=0.106, *q*<.0001, BF_10_=1.8×10^11^); the two caregiver-support dimensions were associated with a smaller SCAN (*r*=-0.062 and -0.057, *q*s<.003; BF_10_=367 and 46), consistent with a buffering influence of supportive caregiving. Across the full adversity-by-network matrix, the SCAN had significant associations with nearly every dimension of adversity; the remaining 14 networks had weaker and sparser effects (Fig. 1e).

To examine which dimension carries this signal, we organized the ten factors into three composites (threat, deprivation, and unpredictability), entered simultaneously into a model for each network along with covariates. Threat was the dominant predictor: a one-standard-deviation increase in threat was associated with a 24% increase in the SCAN’s share of cortex (β=0.0023, *z*=7.30, *p*<.0001; Cohen’s *d*=0.52, BF_10_=1.4×10^11^; Fig. 1g); neither deprivation nor unpredictability remained significantly associated with the SCAN once threat was accounted for (BF_01_=45, *p*=0.40; and BF_01_=66, *p*=0.87). Children at or above one standard deviation of threat devoted, on average, 3.19% of cortex to the SCAN, compared with 2.57% in their least-adversity-exposed peers (Fig. 1g). Among all 15 networks, the SCAN was, by a wide margin, the most adversity-sensitive network in the cortex. The three adversity composites explained nearly seven times more incremental variance in the SCAN’s cortical share (pseudo-ΔR^2^=0.024) than in any other network (Fig. 1c), the closest being the temporal pole (pseudo-ΔR^2^=0.0036), 17 times more than the average of remaining networks and as much as 118 times more than the least affected network. To a first approximation, the imprint of adversity on cortical topography is largely an imprint on the SCAN.

This selectivity held across development and across independent subsamples. The threat-SCAN association persisted two and four years later (2-year: *r*=0.121, *q*<0.001; 4-year: *r*=0.081, *q*=0.004; Fig. 1h), indicating a stable relation across adolescence. To assess whether the prominence of the SCAN was robust to overfitting, we repeatedly split the cohort into independent discovery and replication halves at the family level, so that no genetic or household relationship straddled the divide. Across 20 such partitions, the SCAN had the largest pseudo-ΔR^2^ of all 15 networks (Fig. 1j); there was a significant threat-SCAN association in both halves on each of the 20 splits. The effect was nearly identical across subsamples (discovery β=0.0024; replication β=0.0022), and the rank order of adversity selectivity across networks was highly reproducible between sample halves (*r*=0.94).

A further question is whether this association reflects adversity itself or the stable familial background against which it occurs, including shared genetics, socioeconomic circumstances, neighborhood, and parenting. To address this question, we used the cohort’s embedded sibling structure (456 families with two or more imaged siblings) to estimate the threat-SCAN association within families, holding constant everything siblings share, including most of their adversity exposure, leaving under 1% of the variance in threat within families. Even across these small within-family differences, the sibling with greater threat exposure had the larger SCAN (β=0.0032, *p*=0.039, family fixed-effects with cluster-robust SEs; Fig. 1i); a discordant-pair analysis yielded a consistent, though marginally significant, estimate (β=0.0030, *p*=0.053). This pattern is difficult to reconcile with a between-family confound, which would attenuate once shared influences are held constant, and instead points to an effect that is unusually robust and attributable to threat itself.

### The SCAN Expands by Encroaching on Adjacent Sensorimotor and Cingulo-Opercular Networks

Because cortical surface area is finite, a larger SCAN must come at the expense of other networks, and the selective expansion described above is necessarily a redistribution of territory. Therefore, to examine from where the additional SCAN territory is drawn, we quantified for each network its contribution to the encroaching portion of the SCAN. In the most threat-exposed children (+1SD), this expansion concentrated on a few specific topographical neighbors. On average, the somatomotor dorsal (SMD) network accounted for 34.5% of the encroaching territory, the somatomotor lateral (SML) for 29.3%, and the cingulo-opercular (CO) for 24.8%, with little intrusion elsewhere (Fig. 2b, d). Treating threat as a continuous predictor confirmed this specificity. Each one-standard-deviation increase in threat was associated with greater encroachment of the SCAN into CO (β=0.10), SMD (β=0.09), SML (β=0.08), and AUD (β=0.08) networks (all *p*s<0.001), with no encroachment into higher-order association networks such as frontoparietal, salience, or default-mode (all *p*s>0.53). Thus, with greater threat exposure, the SCAN appears to intrude into the cortex it immediately borders, particularly into sensory and motor systems.

**Figure 2:**
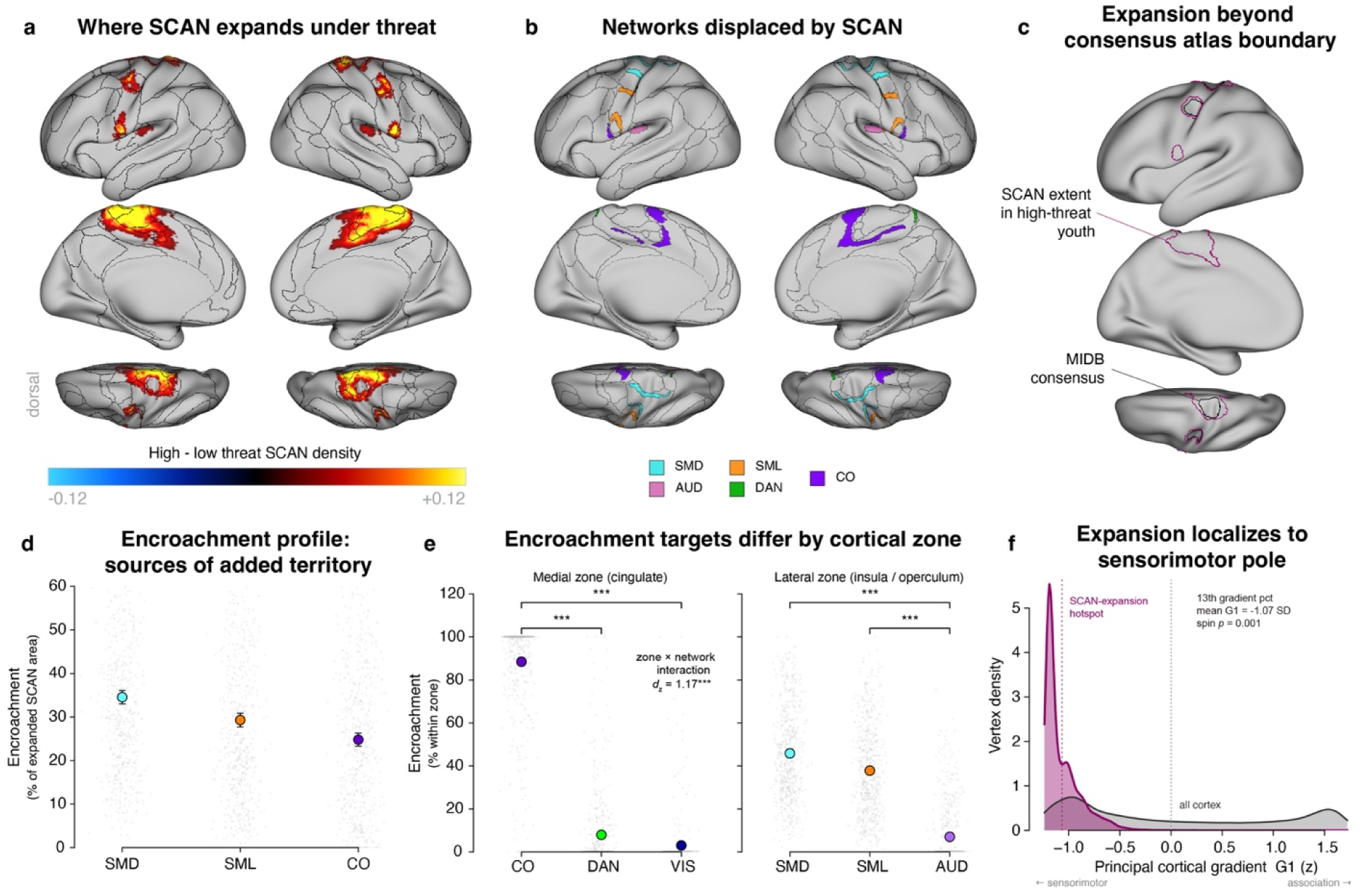
Threat-related SCAN expansion encroaches selectively on sensorimotor cortex. **(a)** Difference in between-subject SCAN density across high- and low-threat youth (high - low, ≥= +1 SD vs ≤ -1 SD), on lateral, medial, and dorsal surfaces; warm vertices mark where SCAN territory expands under threat. **(b)** The networks that the expanded SCAN territory overlaps and displaces, colored by network (SMD, SML, AUD, DAN, CO). **(c)** The SCAN boundary in high-threat youth (magenta) extends beyond the independent MIDB consensus atlas boundary (black; 50% threshold), indicating the expansion is not an artifact of our parcellation. **(d)** Composition of the encroaching territory: of the cortex newly captured by SCAN, the largest shares belong to somatomotor-dorsal (SMD, 34.5%), somatomotor-lateral (SML, 29.3%), and cingulo-opercular (CO, 24.8%) cortex. **(e)** This encroachment is spatially organized; within the medial (cingulate) zone it targets CO almost exclusively (88.5%), whereas within the lateral (insula/operculum) zone it targets SMD (45.8%) and SML (37.8%) – a double dissociation (zone × network interaction, *d*_z_ = 1.17, ***BH-FDR *q* < .001 within zone). **(f)** The expansion hotspot localizes to the sensorimotor pole of the principal cortical gradient (13th percentile, mean G1 = -1.07 SD; spin-test *p* = 0.001), well below the rest of cortex. Points in d-e are individual high-threat youth; error bars are 95% CI. MIDB = Masonic Institute for the Developing Brain.

This encroachment was anatomically organized (Fig. 2e). Along the medial wall it came almost entirely at the expense of the cingulo-opercular network (88.5% of the medial encroaching area), whereas on the lateral surface it displaced the somatomotor networks (45.8% from SMD and 37.8% from SML), with CO scarcely affected laterally (5.4%; Fig. 2b,c).

This crossover was confirmed by a highly significant interaction of network (cingulo-opercular vs somatomotor) and zone (medial vs lateral; *t*(578)=28.2, *p*<.0001, Cohen’s *d*_z_=1.17), with CO preferentially displaced medially (*d*_z_=0.69) and the somatomotor networks laterally (*d*_z_=1.60). This dissociation mirrors the dual anchoring of the SCAN in the cingulate midline and the inter-effector somatomotor nodes, indicating that adversity extends each of the SCAN’s anchors into the territory it abuts.

We then tested whether these displaced networks share a common position in the macroscale functional hierarchy of the cortex, indexed by the principal connectivity gradient running from unimodal sensorimotor to transmodal association cortex. Projecting the vertex-wise map of the adversity-related difference in SCAN extent onto this gradient, we found that this difference was greatest at the sensorimotor end (mean 1.07 SD below the cortical average, the 13th percentile; spin-test *p*=0.001; Fig. 2f). Thus, expansion of the SCAN is oriented down the cortical hierarchy toward sensory and motor systems.

### Adversity Shifts the Functional Coupling of the SCAN from Cognitive-Control Toward Sensorimotor Systems

We next examined whether the expansion of the SCAN is accompanied by a reorganization of its functional connectivity. Using the same individualized parcellations to define network nodes, we computed the resting-state functional connectivity between the SCAN and every other network and tested its association with threat. Seven of the fourteen SCAN connections were significant after correction, in a coherent pattern (Fig. 3d-e). Greater threat was associated with stronger coupling of the SCAN with sensory and motor systems, most strongly with the dorsal somatomotor network (β=0.034, q<.0001) and more modestly with the auditory network (β=0.018, *q*<.0001) and the parieto-occipital network (β=0.016, *q*<0.0001; Fig 3a,c,d). Over the same range of exposure, the coupling of the SCAN with the association and control systems that ordinarily regulate it grew weaker, including the salience (β=-0.016, *q*<.0001), frontoparietal (β=-0.016, *q*<0.0001), cingulo-opercular (β=-0.010, *q*=0.014), and the parietal memory (β=-0.010, *q*=0.008) networks. Thus, consistent with its structural expansion, adversity shifts the functional coupling of the SCAN away from cognitive-control systems and toward sensorimotor cortex.

**Figure 3:**
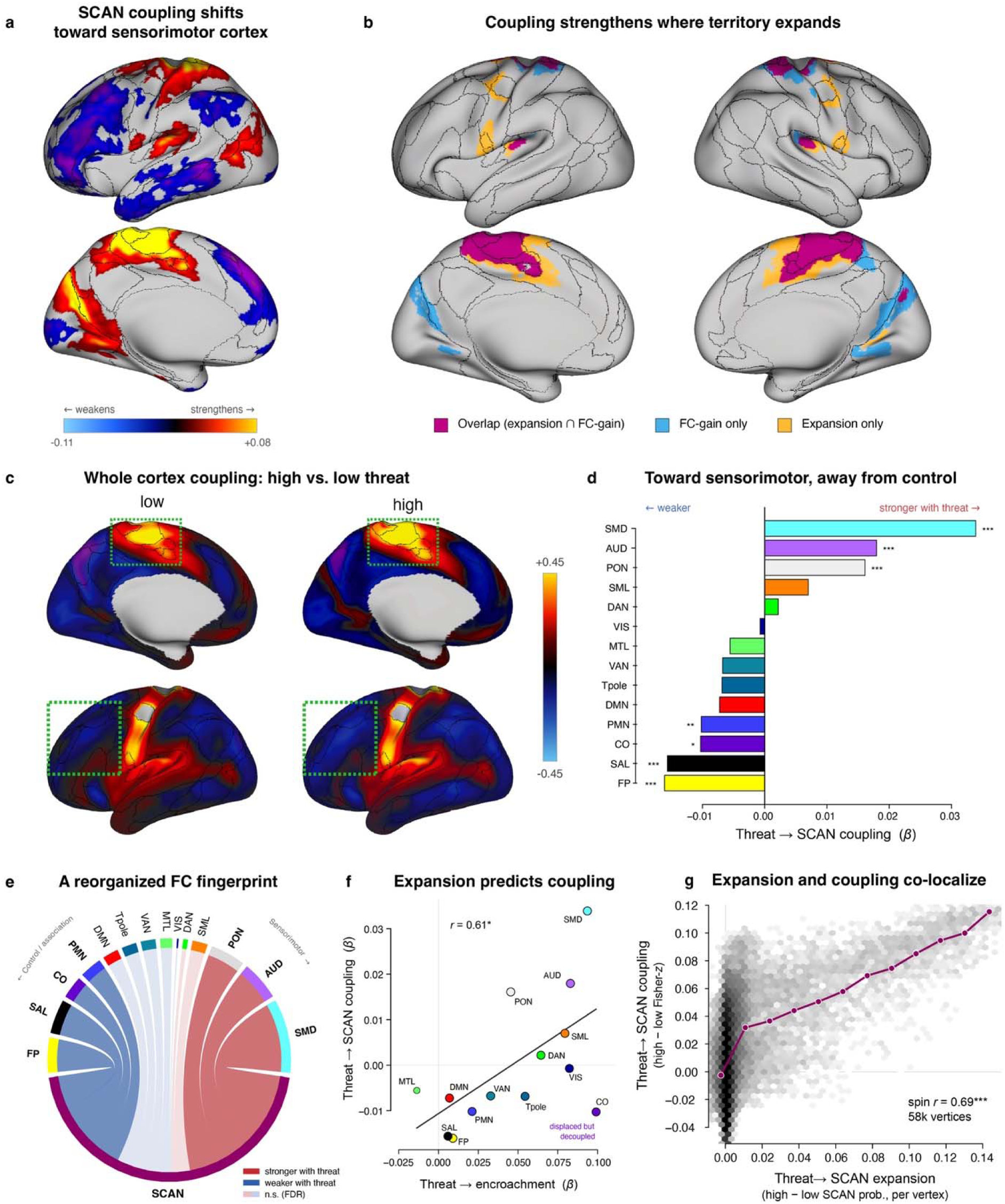
Threat shifts SCAN’s functional coupling toward sensorimotor cortex, mirroring spatial expansion. **(a)** Vertexwise change in SCAN-seed functional connectivity between high- (+1SD) and low- (-1SD) threat youth (high - low). SCAN coupling strengthens with sensorimotor/perisylvian cortex (warm) and weakens with association/control cortex (cool). **(b)** Conjunction of the two maps: cortex showing both SCAN territorial expansion and a coupling gain (overlap, magenta) versus FC-gain only (blue) or expansion only (orange); the magenta overlap marks where structure and function change together. **(c)** Group-mean SCAN-seed connectivity in low- versus high-threat youth. **(d)** Threat → SCAN coupling effects (β) across all 14 partner networks: coupling strengthens most with somatomotor-dorsal (SMD, +0.034), auditory (AUD), and parieto-occipital (PON) cortex, and weakens with salience (SAL, -0.016), frontoparietal (FP), cingulo-opercular (CO) control, and parietal memory (PMN) networks (7/14 FDR-significant; \**q* < .05, \*\**q* < .01, \*\*\**q* < .001). **(e)** The same reorganization as a connectivity fingerprint: SCAN’s couplings strengthen toward sensorimotor partners (red) and weaken toward control partners (blue). **(f)** Network-level structure-function correspondence: networks that SCAN encroaches on more under threat also strengthen their coupling (*r* = 0.61, permutation *p* = 0.020, n = 14 networks); CO is the exception – heavily displaced but functionally decoupled. **(g)** The same correspondence at the vertex level is stronger and highly reliable (spin-test *r* = 0.69, *p* = 0.001, ∼58,000 vertices): where SCAN expands is where its coupling increases.

The networks toward which the connectivity of the SCAN shifted were largely those into which it expanded structurally (Fig. 3b), suggesting that the territorial and connectivity differences are two expressions of one reorganization. Across all 14 networks, the magnitude of threat-related encroachment predicted the magnitude of threat-related connectivity change, such that the more a network’s territory was annexed by the SCAN under threat, the stronger its coupling with the SCAN (*r*=0.61, permutation *p*=0.020; Fig. 3f). A vertexwise analysis seeded from a fixed group-template SCAN reproduced this correspondence at a much higher resolution.

Across the cortical surface, the threat-related difference in SCAN connectivity tracked the threat-related difference in SCAN territory vertex by vertex (spin-test *r*=0.69, *p*=0.001, Fig. 3g), such that the SCAN strengthened its coupling with the cortex into which it expanded. This connectivity increase peaked at the sensorimotor pole of the principal gradient (13.0^th^ percentile), coinciding with the territory that the SCAN annexed (13.3^th^ percentile; Fig. 2). Thus, topographical annexation and functional coupling co-vary across networks, consistent with a single process in which the enlarged sensorimotor territory of the SCAN is accompanied by, and may contribute to, its strengthened sensorimotor connectivity.

One notable exception to this pattern, however, was the cingulo-opercular network. Whereas this area ceded the most medial territory to the SCAN, its coupling with the SCAN was *weaker* in youth who were exposed to greater threat (Fig. 3f), opposite to the sensorimotor networks that the SCAN annexed. Therefore, the SCAN appears to absorb sensorimotor cortex into its functional fold while supplanting the CO control network it overruns. This difference between the systems an expanded SCAN integrates versus those it displaces motivated us to investigate the implications of an enlarged and sensorimotor-shifted SCAN for cognition and behavior.

### SCAN Expansion Exacts a Developmental Cost to Cognition

To assess the functional consequence of this reorganization, we tested whether the topography of the SCAN assessed at baseline mediates the association between early threat and crystallized and fluid intelligence six years later, bootstrapping the indirect effect over 5,000 family-clustered resamples. We found that greater threat exposure was associated with a larger SCAN (a-path β=0.0023, *p*<0.001; Fig. 4a), which in turn predicted lower crystallized intelligence (indirect=-0.21, 95% CI [-0.42, -0.03], bootstrap *p*=0.016); the fluid pathway was not significant (indirect=-0.18, 95% CI [-0.44, 0.05]). This crystallized mediation recurred at every testable wave (year-2=-0.22, *p*<0.001; year-4=-0.35, *p*=0.023; year-6=-0.21, *p*=0.016; Fig. 4b), suggesting that the cost of a larger SCAN was sustained across adolescence. These associations also held out of sample; in cross-validated models, the baseline SCAN forecast year-6 crystallized cognition in held-out children (Fig. 4d), improving prediction beyond demographic covariates (CV ΔR^2^=0.011, permutation *p*=0.001, N=2,524). The incremental value of the SCAN beyond threat itself was modest (CV ΔR²= 0.003), which was expected given that the SCAN was on the causal path between threat and cognition.

**Figure 4:**
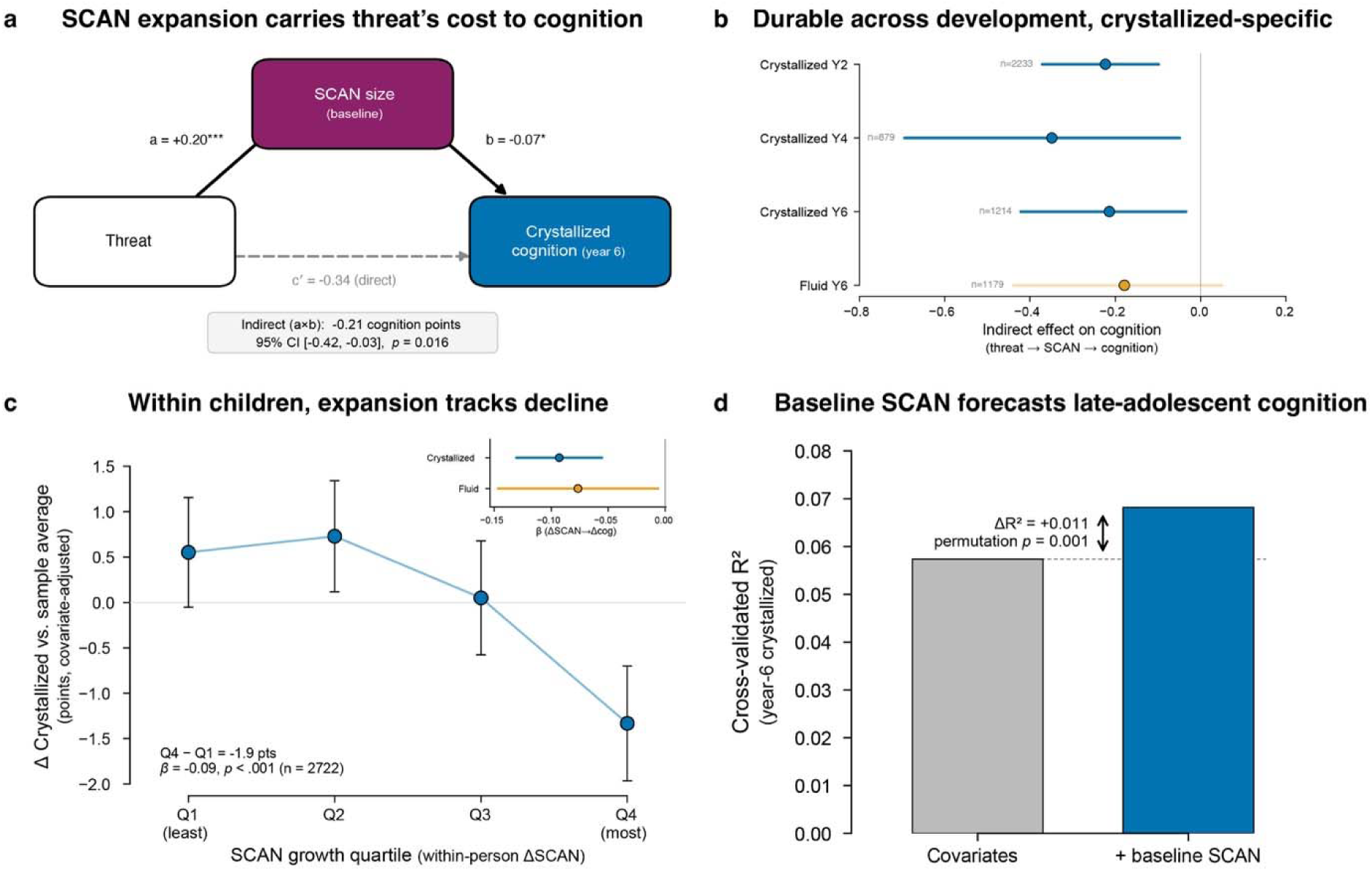
Threat-driven SCAN expansion carries a cognitive cost. **(a)** Mediation model: threat predicts greater SCAN expansion (a = +0.20), which in turn predicts lower year-6 crystallized cognition (b = −0.07); the indirect path is -0.21 cognition points (95% CI [-0.42, - 0.03], bootstrap *p* = 0.016, n = 1214), with the direct threat → cognition path (c′ = -0.34) shown faint. Standardized coefficients on arrows; the indirect effect is in raw cognition points. **(b)** The indirect effect replicates across development – crystallized cognition at Year 2 (n = 2233), Year 4 (n = 879), and Year 6 (n = 1214), all with CIs excluding zero – and is crystallized-specific; the year-6 fluid effect (orange) crosses zero. **(c)** The relationship holds within children over time; youth in the highest within-person SCAN-growth quartile (Q4) show the steepest crystallized decline relative to the sample average (Q4 - Q1 = -1.9 points; all-waves mixed-model β = -0.09, *p* < .001, n = 3,879 observations from 2,722 children). Inset: within-person ΔSCAN → Δcognition is significant for both crystallized (blue) and fluid (orange). **(d)** Out-of-sample prediction: adding baseline SCAN to a covariate model improves cross-validated prediction of year-6 crystallized cognition (ΔR^2^ = +0.011, permutation *p* = 0.001). Color encodes cognitive domain (blue = crystallized, orange = fluid); the SCAN node is magenta. Error bars are 95% CI.

Although these analyses relate a baseline brain measure to later cognition, they leave open the question of whether cognitive change *within a child* tracks with the threat-related expansion of the SCAN. In within-person models across all waves, greater threat predicted continued enlargement of the SCAN beyond each child’s own baseline (β=0.042, *p*=0.002); further, this growth predicted declining cognition over the same period for both crystallized (β=-0.093, *q*<0.001) and fluid ability (β=-0.076, *q*=0.033; Fig. 4c). Threat itself independently predicted crystallized decline (β=-0.118, *p*<0.001). Notably, fluid ability tracked within-person SCAN growth despite showing no cross-sectional mediation, extending the longitudinal cost to both domains. Thus, cross-sectional and within-person analyses suggest that children whose SCAN expanded most following adversity show the steepest declines in cognition.

Finally, with respect to psychopathology, no CBCL subscale showed a significant indirect effect of threat through SCAN topography (none survived FDR, *q*s>0.8; all bootstrap CIs included zero), and within-person SCAN change did not predict change in any subscale (qs>0.6). Because the cognitive pathway is supported by converging cross-sectional, within-person, and out-of-sample evidence, we interpret it as a likely developmental consequence of SCAN expansion; by contrast, we found no evidence that SCAN expansion relates to broadband psychopathology (see full CBCL results in the Supplement).

## Discussion

How the early environment becomes durably inscribed in the organization of the developing cortex is a central question in developmental neuroscience. Here we examined whether, and along which dimensions, early adversity reshapes individualized cortical network topography across adolescence, and with what functional and behavioral consequences. An unbiased screen across 15 networks and 10 domains of adversity in a large sample of youth converged on a single result: threat, but not deprivation or unpredictability, was associated with pronounced expansion of the SCAN, an effect nearly seven times larger than that for the next-most-affected network. This expansion was geometrically organized, encroaching on adjacent somatomotor and cingulo-opercular cortex and localizing to the sensorimotor pole of the cortex’s sensorimotor-association axis^22,34,35^. The same expansion was accompanied by a coordinated shift in the SCAN’s functional coupling toward sensorimotor and away from control networks; further, it carried a cognitive cost that replicated across development, persisted within individual children over time, and predicted cognition in held-out youth.

The function of the SCAN, identified by Gordon and colleagues (2023)^24^, may explain why it appears to expand as a function of exposure to threat in particular. The SCAN is a system of inter-effector regions that interrupt the motor homunculus and, rather than controlling isolated movements, integrate goal-directed action with whole-body physiological control. The network connects densely with the cingulo-opercular network and with subcortical and visceral effectors, including autonomic structures such as the adrenal medulla^36^, and has been implicated in arousal, autonomic regulation, action readiness, and the coupling of bodily state to behavior.

Because chronic threat places sustained demand on physiological mobilization, it may repeatedly engage the SCAN during a developmental window in which cortical territory is allocated competitively among networks^22^. If cortical territory is shaped by how often a network is engaged, with more active regions expanding into less-active neighbors through use-dependent plasticity^37^, then a childhood spent repeatedly mobilizing for action should enlarge the SCAN at the expense of the neighbors with which it competes for growth. In this sense, threat-related SCAN expansion may represent the topographic consequence of an action system that has been chronically engaged, a developmental form of biological embedding in which a history of threat is written into the cortical territory occupied by the network. It is notable that this imprint falls on a system built for physiological mobilization, given that early threat is studied most often in the context of fear-learning, salience, and frontolimbic circuits. Because our analysis was restricted to cortical networks, it does not address the subcortical structures, such as the amygdala, that are central to classical threat circuitry. Within the cortex, however, the strongest imprint of threat fell not on prefrontal or salience systems but on a network whose defining role is to couple the brain to the body’s physiological state.

In more threat-exposed youth, expansion of the SCAN fell almost entirely on its immediate sensorimotor and cingulo-opercular neighbors, leaving higher-order association networks such as frontoparietal, salience, and default mode untouched. This expansion was spatially patterned, extending into the cingulo-opercular network along the midline but into somatomotor networks on the lateral surface, a double dissociation between cortical location and the functional network it displaced. Because the SCAN is itself anchored in both the cingulate midline and the lateral inter-effector nodes of the motor strip, each of its anchors appears to have extended into the cortex it directly abuts. Thus, adversity appears to amplify rather than distort the network’s form, extending the SCAN along its existing axes. The result is a developing cortex that, under threat, gives more of its surface over to an action system and less to the neighbors it overtakes.

The territorial expansion of the SCAN was accompanied by a parallel shift in its functional coupling, which strengthened toward the same sensorimotor network upon which its territory had encroached. These structural and functional changes were spatially aligned, with the extent of the SCAN’s encroachment predicting how much its functional coupling with a given system changed, an alignment evident even at the individual-vertex level. Therefore, our finding that the enlarging territory and strengthening coupling of the SCAN emerge together suggests that the topographical and functional changes reflect a single reorganization rather than two separate processes. An important exception to this finding was that although the cingulo-opercular cortex ceded more medial territory to the SCAN than did any other network, its coupling with the SCAN was weaker rather than stronger in more threat-exposed youth, differentiating it from the somatomotor networks that SCAN both annexed and to which it became more tightly coupled. This distinction tracks the functional roles of the two systems; the somatomotor cortex shares the orientation of the SCAN toward action and is drawn into its functional fold, whereas the cingulo-opercular network is a control system whose territory the SCAN overtakes without assimilating its regulatory function. In effect, therefore, the SCAN integrates the sensorimotor cortex it absorbs while supplanting the control cortex it displaces.

In this context, the reorganization of the SCAN had a measurable cost to cognitive functioning. Expansion of the SCAN mediated the association between early threat and poorer cognitive performance, an effect that was most consistent for crystallized ability and was found at every assessment wave. The same pathway was evident out-of-sample, with baseline SCAN size forecasting later crystallized cognition in youth the models had not seen, and within individuals over time (adolescents whose SCAN expanded most over time had the steepest declines in cognitive functioning). The concentration of this effect in crystallized ability is noteworthy. Because crystallized knowledge accumulates gradually across development, a cortex that is progressively tuned toward action may accrue such knowledge more slowly, leading to the emergence of a durable deficit^38^. In contrast, fluid ability followed the pace of SCAN growth within children rather than reflecting this cumulative gap. Because the same reorganization that enlarges the SCAN also weakens its coupling with the control systems that support higher cognition, this deficit may be the behavioral expression of that decoupling.

This cognitive cost also bears on the dimensional framework that motivated our screen, which has classically linked cognitive deficits to deprivation rather than to threat^25,27^. Rather than disconfirming that mapping, our findings reveal a second and previously unmapped route to the same endpoint, by which threat affects cognitive functioning by reorganizing an action system rather than by degrading associative input attributed to deprivation. Adaptation-based accounts of development offer a way to interpret this cost, proposing that early adversity calibrates the developing brain to the demands it faces and that the resulting specialization carries tradeoffs whose costs fall on abilities that are less relevant to a child’s specific environment^39,40^. The threat-specific effect matches the prediction of this account that distinct dimensions of adversity shape distinct abilities. Therefore, an enlarged, sensorimotor-shifted SCAN may reflect a cortex organized for action in response to threat, an adjustment that could be adaptive for a child in a dangerous environment even as it exacts a cost on more abstract cognitive abilities. We advance this interpretation cautiously: while our data establish that greater SCAN expansion accompanies lower cognition, we cannot establish a causal link.

Together, these findings recast how we might best understand the imprint of the environment on the developing brain. Adversity has often been treated as a diffuse influence that is spread across many neural systems. Our results suggest, in contrast, that its imprint on cortical network organization is concentrated in a single network whose expansion, connectivity, and cognitive cost cohere into one developmental process. In a recent examination of hundreds of environmental exposures, socioeconomic factors were the strongest correlates of functional connectivity and cortical thickness, with effects concentrated in sensory and motor cortex and overlapping with arousal-related neuromodulatory systems^10^. Threat-related SCAN expansion supplies a candidate mechanism for that concentration. Because our effect held for threat specifically, survived adjustment for socioeconomic factors, and appeared within individual children, the shared sensorimotor localization is unlikely to mean that threat is simply a proxy for socioeconomic disadvantage. More plausibly, threat and socioeconomic adversity act as distinct exposures that converge on the same parts of cortex.

We should note four limitations of this study. First, our design is observational and the central association between threat and SCAN expansion is cross-sectional. Although the cognitive pathway draws on longitudinal and within-person evidence, we cannot determine whether a larger SCAN precedes or follows threat exposure, or exclude the possibility that the SCAN expansion and cognition share common antecedents rather than having a causal relation. Second, the link between SCAN expansion and cognition could reflect residual socioeconomic confounding. We addressed this possibility by using closely matched covariates and within-person models that hold constant between-child differences, although we recognize that such strategies reduce rather than eliminate this concern. Third, our parcellation assigns each cortical location to a single network in a winner-take-all fashion, whereas the underlying connectivity is likely more graded than this hard boundary implies; relatedly, the boundary between the SCAN and the adjacent salience and cingulo-opercular networks depends on the parcellation framework, and the apportionment of encroached territory should be interpreted with that dependence in mind. Finally, the analytic sample, while large, differed modestly from the rest of the cohort in sociodemographic characteristics, which may bound generalizability.

Despite these limitations, our findings are important in documenting that early threat leaves a specific and legible mark on the topographic organization of the developing brain. The effect is concentrated on a single functionally coherent system rather than altering the overall cortical map across networks. The targeted reorganization that we documented in this study is what dimensional models of adversity have long anticipated but seldom localized: a developing cortex that, under threat, is biased toward action and arousal at the expense of the association systems on which higher cognition depends. Our findings that this neural signature is identifiable in individual children, from a single scan, and at population scale, helps to elucidate how exposure to threat can become embedded in the architecture of the developing brain.

## Methods

### Participants

Participants were drawn from the Adolescent Brain Cognitive Development (ABCD) Study, a longitudinal cohort study of 11,880 youth enrolled at ages 9-10 years (*M*=9.98, *SD*=0.63) across 21 research sites in the United States.^41^ For full enrollment and inclusion/exclusion criteria, see ABCD Release 7.0 documentation at https://abcdstudy.org/. Children were assessed at four waves: baseline (*M*=9.98 years), 2-year follow-up (*M*=12.03 years), 4-year follow-up (*M*=14.20 years), and 6-year follow-up (*M*=16.10 years). To be included at a given wave, participants needed to have usable resting-state fMRI data (see below) and complete early life adversity (ELA) data. After applying these criteria, analytic samples were N=4,525 at baseline; N=4,347 at 2-year follow-up; N=3,983 at 4-year follow-up; and N=2,687 at 6-year follow-up. Families with multiple enrolled siblings (N=456) were retained and the resulting non-independence was handled statistically throughout (see Data Analyses). Mediation analyses linking ELA exposure to behavioral outcomes were conducted with participants who had complete baseline imaging, ELA, and 6-year outcome data (NIH Toolbox Fluid Cognition: N=1,179; NIH Toolbox Crystallized Cognition: N=1,214; CBCL subscales: N=1,513). All participants provided written assent, and written informed consent was obtained from a parent or legal guardian; all procedures were approved by both a central institutional review board at the University of California, San Diego and each site’s IRB.

### Resting State fMRI Acquisition and Preprocessing

Resting-state fMRI data were obtained from the ABCD BIDS Community Collection (ABCC). Data were preprocessed using the ABCD-BIDS pipeline^42^, a modified version of the Human Connectome Project processing pipeline. Motion censoring was applied using a framewise displacement threshold of FD<0.2 mm, and only participants with a minimum of 10 minutes of low-motion data remaining post-censoring were retained; for participants who exceeded this threshold, frames were randomly sampled to yield exactly 10 minutes of data for connectivity matrix generation. Full preprocessing procedures are described in Hermosillo et al. (2024)^43^.

### Functional Network Topography

Individual-specific functional network topography was quantified using the template matching (TM) algorithm described in Hermosillo et al. (2024)^43^. Network templates were derived from a normative subsample of ABCD participants (n=164, ages 9-10 years) using seed-based correlations thresholded at z≥1 (top 15.9% of connections). For each participant, connectivity matrices were z-scored separately for each cortical hemisphere and subcortical region to normalize connectivity across regions with differing signal-to-noise characteristics; the resulting whole-brain connectivity pattern at each grayordinate was then compared to each network template using eta-squared (η²) as the similarity metric, and each grayordinate was assigned to its best-matching network. This procedure was applied across all 91,282 cortical and subcortical grayordinates in the CIFTI framework. We then restricted our analyses to the 59,412 cortical surface grayordinates, excluding subcortical structures.

Fifteen bilateral functional networks were identified using the ABCD Template Matching V2 atlas (Masonic Institute for the Developing Brain; midbatlas.io), including the Somato-Cognitive Action Network (SCAN^24^) along with 14 previously characterized networks: Default Mode (DMN), Visual (VIS), Frontoparietal (FP), Dorsal Attention (DAN), Ventral Attention (VAN), Salience (SAL), Cingulo-Opercular (CO), Somatomotor Dorsal (SMD), Somatomotor Lateral (SML), Auditory (AUD), Temporal Pole (Tpole), Medial Temporal Lobe (MTL), Parietal Memory (PMN), and Parieto-Occipital (PON). Network topography was operationalized as the proportion of total cortical surface area (mm^2^) assigned to each network, capturing each network’s relative spatial extent within a given individual.

### Early Life Adversity

Ten dimensions of early life adversity (ELA) were drawn directly from Brieant et al.^44^, who derived them from the ABCD Study baseline assessment; we adopted their published factor solution and factor scores without modification. Briefly, Brieant and colleagues applied exploratory factor analysis (EFA) with oblique rotation to 60 environmental and experiential items spanning trauma exposure, caregiver psychopathology, parenting, family conflict, and family- and neighborhood-level socioeconomic disadvantage, drawn from parent- and youth-report instruments as well as residential address records. Their ten-factor solution was selected on the basis of model fit and interpretability, retaining the most parsimonious solution that reached excellent fit (RMSEA=0.02, CFI=0.97, TLI=0.96; thresholds RMSEA<0.05, CFI and TLI>0.95); solutions with additional factors were rejected because they contained factors with no meaningful item loadings. The ten factors captured: (1) caregiver psychopathology; (2) socioeconomic disadvantage and lack of neighborhood safety; (3) secondary caregiver lack of support; (4) primary caregiver lack of support; (5) youth-reported family conflict; (6) caregiver substance use and biological caregiver separation; (7) family anger and arguments; (8) family verbal and physical aggression; (9) physical trauma exposure; and (10) caregiver lack of supervision. Following Brieant et al., each participant’s score on a given factor was computed as a weighted average of their item responses, with each item weighted by its factor loading, and the resulting scores were standardized (z-scored) prior to analysis.

Because the ten factors are relatively granular and intercorrelated (Supplementary Figure S1) – for example, primary and secondary caregiver support load on separate factors – we performed a further theory-guided reduction, grouping the ten dimensions *a priori* into three composites following the threat, deprivation, and unpredictability taxonomy^27,26,45^. This step improves interpretability and reduces collinearity among predictors while preserving the established dimensional structure of adversity. A threat composite was constructed as the mean of four ELA dimensions reflecting exposure to direct interpersonal threat: physical trauma exposure; family verbal and physical aggression; youth-reported family conflict; and family anger and arguments. A deprivation composite was constructed as the mean of socioeconomic disadvantage/neighborhood unsafety, primary caregiver lack of support, secondary caregiver lack of support, and caregiver lack of supervision. An unpredictability composite was constructed as the mean of caregiver psychopathology and caregiver substance use and separation. All three composites were z-scored after averaging. For the three composites, all factors were scored such that higher values reflect higher adversity. ELA factor and composite intercorrelations are presented in Supplementary Figure S1.

### Behavioral Outcomes

Cognitive ability was indexed using NIH Toolbox Cognition Battery^46^, a standardized, computerized battery normed to a nationally representative U.S. sample. The fluid cognition composite aggregates performance across five tasks assessing processing speed, sustained attention, working memory, cognitive flexibility, and episodic memory. The crystallized cognition composite reflects accumulated verbal knowledge, derived from vocabulary comprehension and oral reading recognition subtests. Both composites are expressed as age-corrected standard scores (*M*=100, *SD*=15 in the normative sample) and were assessed at the 6-year follow-up.

In addition, we assessed mental health using the parent-reported Child Behavior Checklist (CBCL^47^), a well-validated broadband measure of emotional and behavioral problems in youth ages 6-18 years. Fourteen CBCL subscales assessed at the 6-year follow-up (ages 15-18 years) were examined as outcomes in mediation analyses including the three broadband scales (Total Problems, Internalizing, Externalizing), eight narrow-band syndrome scales (Anxious/Depressed, Withdrawn/Depressed, Somatic Complaints, Social Problems, Thought Problems, Attention Problems, Rule-Breaking, Aggressive Behavior), and three DSM-5-oriented scales (Depressive Problems, ADHD, Conduct Problems). The CBCL has been shown to have good internal consistency (α=0.75-0.84) and test-retest reliability (*r*=0.78-0.88)^48^. Benjamini-Hochberg false discovery rate (FDR) correction was applied within each outcome domain, correcting the two NIH Toolbox cognition measures and the 14 CBCL subscales separately; component a- and b-paths are reported uncorrected as constituents of each indirect effect. To isolate prospective changes in symptom burden rather than pre-existing psychopathology, each participant’s baseline score on the matching CBCL subscale was included as a covariate in the corresponding CBCL mediation model.

### Data Analyses

We first screened all ten ELA dimensions against the topography (proportion of total cortical area) of all fifteen networks to identify which associations were robust *(bivariate analyses)*, then estimated the unique contribution of each adversity dimension while controlling for the others *(multivariate composite models)*. Having localized the effect to threat and the SCAN, we characterized how the expanded SCAN redistributes cortical territory *(encroachment analysis)* and how this territorial difference is reflected in functional connectivity *(functional connectivity analysis)*. Finally, we tested whether SCAN topography links threat to cognitive and psychiatric outcomes in late adolescence *(mediation analyses)*, and whether these associations hold within individual children over time *(within-person longitudinal analyses)*.

#### Bivariate ELA-Network Associations

To characterize the broad associations between ELA dimensions and functional network topography, we first fit ordinary least squares (OLS) regression models regressing each network proportion on each ELA factor individually, covarying for age at scan, sex, and mean framewise displacement. Study site was entered as a fixed effect (dummy-coded) to account for scanner and protocol differences across the 21 ABCD sites; because study site fully determines scanner manufacturer and serial number in ABCD (each nested within site), these fixed-effect indicators absorb between-scanner as well as between-site variance. Sibling non-independence was accommodated with family-cluster-robust (sandwich) standard errors rather than a family random intercept, whose variance was negligible in this sample (only 11.2% of families contributed more than one imaged participant, yielding a singular random-effects covariance). Because these models are estimated by OLS, they involve no iterative random-effects fitting; the alternative random-effects specifications we examined as robustness checks (random site, random family, their combination, and family nested within site) all converged and left the associations essentially unchanged (Supplemental Table S1). We did not covary for socioeconomic status because socioeconomic disadvantage constitutes one of the ELA dimensions under study (a component of the deprivation composite), and adjusting for it would remove variance attributable to adversity itself. Partial correlations were derived from the model cluster-robust test statistics. To test the stability of observed effects, this analysis was repeated separately at each wave. FDR correction (Benjamini-Hochberg) was applied within wave across all 150 tests (10 ELA dimensions x 15 networks). An analogous set of models was fit using the three ELA composites, with FDR applied across 45 tests (3 composites x 15 networks) per wave.

#### Multivariate Composite Models

To estimate the unique contribution of each adversity dimension while controlling for shared variance among dimensions, we fit a single model per network entering the three ELA composites (threat, deprivation, and unpredictability; see above) simultaneously along with age, sex, and mean FD. As in the bivariate analyses, these were OLS models with study site entered as a fixed-effect indicator and family-cluster-robust standard errors to account for sibling non-independence. FDR correction was applied across all 45 composite-network combinations per wave. To quantify each network’s sensitivity to ELA, we computed a pseudo-ΔR^2^, the increase in variance explained (R2) when the three composites were added to a covariate-only OLS model (pseudo-ΔR^2^ = R^2^_full_– R^2^_covariate-only_).

#### Robustness and Replication Analyses

We evaluated the robustness of the threat-SCAN association in three ways. First, to assess stability across development, we refit the bivariate models at each follow-up wave. Second, to guard against overfitting, we repeatedly partitioned the baseline sample into independent discovery and replication halves at the family level, so that no family was split across halves; across 20 such partitions, we compared the variance explained by adversity in each network’s size (pseudo-ΔR^2^) and tested the association between adversity and SCAN topography separately within each half. Third, to distinguish the contribution of threat from the stable familial background against which it occurs, we leveraged the embedded sibling structure of the ABCD data (456 families with two or more imaged siblings), estimating the threat-SCAN association within families via family fixed-effects models with cluster-robust standard errors and a complementary discordant-sibling-pair analysis.

#### Encroachment Analysis

Following the multivariate models, we wanted to determine which networks the SCAN displaces as it expands with greater threat exposure and whether this encroachment is spatially organized according to the anatomical footprint of the SCAN. Because these analyses identified threat as the primary adversity dimension associated with SCAN territory, threat served as the predictor of interest here and in all subsequent analyses. We characterized encroachment in two complementary ways. First, to establish the source of additional territory of the SCAN, we isolated the encroaching portion of each participant’s SCAN, comprising vertices that were labeled SCAN in that participant’s parcellation but were assigned to a different, non-SCAN network in the group-level template. We then computed an encroachment profile, defined as the proportion of this territory assigned to each non-SCAN network in the group template. Profiles were computed in high-threat youth (threat ≥+1 SD, N=579) separately for medial cortex (|x|<20 mm; encompassing cingulate and medial prefrontal regions) and lateral cortex (|x|≥20 mm; encompassing insula, opercular, and lateral prefrontal regions), and were compared across networks using FDR-corrected pairwise *t*-tests within each anatomical zone.

Second, to assess whether encroachment onto a given network scaled continuously with adversity, we computed, for each non-SCAN network, the fraction of that network’s group-template territory relabeled SCAN in a participant’s parcellation. We regressed this displaced fraction on the threat composite via OLS, with study site entered as a fixed-effect indicator and family-cluster-robust standard errors (as in the primary analyses), adjusting for age, sex, and mean FD, with BH-FDR applied across networks, and formally tested the medial-lateral dissociation using a network (cingulo-opercular vs somatomotor) by zone (medial vs lateral) interaction contrast. Finally, to situate the expansion of the SCAN within the macroscale organization of cortex, we projected the vertexwise map of the high-versus-low threat difference in SCAN extent onto the principal cortical connectivity gradient, which runs from unimodal sensorimotor to transmodal association (S-A) cortex^34^. We then tested whether the vertices of greatest SCAN expansion occupied a non-random position along this gradient by comparing their mean gradient value against a null distribution of 1,000 spin-test rotations.

#### Functional Connectivity

Resting-state data were drawn from the ABCC as XCP-D-postprocessed dense timeseries^49^ in fsLR-32k CIFTI grayordinate space. Briefly, data as downloaded were denoised using XCP-D’s ABCD mode, comprising 36-parameter confound regression (six motion parameters and the mean white-matter, cerebrospinal-fluid, and global signals, together with their temporal derivatives and quadratic terms), 3dDespike, interpolation of high-motion volumes, and a 0.01-0.08 Hz band-pass filter, and were spatially smoothed on the surface with a 6 mm full-width-at-half-maximum kernel. We further censored frames at FD≥0.2 mm and retained participants with at least 375 low-motion frames (5 minutes at TR=0.8 s) for FC analyses. For each participant, we averaged the surviving frames within each of the 15 networks defined by that participant’s individualized parcellation, computed the pairwise Pearson correlations among the network timeseries, and Fisher z-transformed them.

Associations between each of the 14 SCAN-to-network FC values and the threat composite were tested via OLS with study site entered as fixed-effect indicators and family-cluster-robust standard errors (as in the primary analyses), covarying for age, sex, mean FD, and number of retained frames. BH-FDR correction was applied across the 14 SCAN-specific pairs. To resolve this association at the level of individual vertices, we also computed vertexwise seed-based connectivity using a fixed group-template SCAN seed (1,336 vertices from the MIDB probabilistic atlas^43^), identical across participants so that the seed did not depend on each child’s individualized parcellation. For each participant, we estimated the correlation between the mean SCAN timeseries and every cortical vertex (frames with FD<0.2 mm) and Fisher z-transformed these values. We formed group-mean seed-connectivity maps for high- (≥+1 SD threat; n=571) and low-threat (≤-1 SD; n=550) youth and took their difference (high-low) as the threat-related difference in SCAN coupling at each vertex; these subsamples are smaller than in the encroachment analysis because the connectivity analysis additionally required usable functional connectivity data. This map was compared against the vertexwise SCAN-expansion (encroachment) difference map by spatial correlation, with significance assessed against 1,000 spin-test rotations, and the gradient position of its peak vertices along the S-A axis was quantified as in the encroachment analysis.

#### Associations with Behavior in Late Adolescence

Mediation analyses tested whether the association between threat exposure (assessed at baseline but retrospective in nature) and behavioral outcomes at the 6-year follow-up was statistically mediated by baseline SCAN proportion using a two-path model (threat → SCAN → outcome). Path a was estimated via OLS with study site as fixed-effect indicators and family-cluster-robust standard errors, covarying for baseline age, sex, and mean FD. Paths b and c′ were estimated simultaneously via OLS with the same study-site fixed effects and family-cluster-robust standard errors (matching path-a and the primary analyses), covarying for sex and 6-year follow-up age, with the matching baseline CBCL subscale added as a covariate for CBCL outcomes to isolate prospective symptom change. The indirect effect (β_a_ x β_b_) was tested via 5,000-iteration cluster bootstrap resampling families with replacement; significance was defined as the 95% percentile CI excluding zero.

BH-FDR was applied within each outcome domain, correcting the two cognitive outcomes and the 14 CBCL subscales separately. Because effects were largely specific to cognition, we next assessed the temporal robustness of the mediation, re-estimating it with crystallized scores from each available follow-up (years 2, 4, and 6). Fluid cognition was administered only at baseline and year-6 and could therefore be tested only at year-6.

Finally, to test whether these associations reflect developmental changes within individual children rather than stable between-person differences, we conducted within-person longitudinal analyses examining change scores relative to each participant’s first available wave, pooling all follow-up observations across waves via MixedLM with a random intercept per participant and cluster-robust sandwich standard errors clustered by family. We first tested whether ELA predicted subsequent growth of the SCAN controlling for each participant’s own baseline SCAN proportion, change in mean FD, years elapsed, age at baseline, and sex (N=6,129 observations; 3,454 participants). We next tested whether SCAN growth, in turn, predicted change in crystallized and fluid cognition controlling for baseline cognition, baseline SCAN, change in mean FD, years elapsed, baseline age, and sex, with BH-FDR applied across the two cognitive outcomes (crystallized N=3,879 observations; fluid N=1,032); and whether ELA independently predicted change in crystallized cognition with the same covariate set (N=3,879 observations). The same within-person change models were applied to all 14 CBCL subscales, with BH-FDR applied across the 14 subscales (N=6,094 observations from 3,442 children), to test whether SCAN change related to within-person change in psychopathology.

## Data Availability

Data from the ABCD Study, including derivatives processed as part of ABCC, are available through the NIH Brain Development Cohorts (NBDC) Data Hub (https://www.nbdc-datahub.org/) and require a Data Use Agreement for access. Analyses used ABCD Release 7.0.

## Code Availability

All code used for the analyses described in this manuscript are available in an online repository (https://github.com/cantonacci/adversity_topography). Software packages incorporated into the analysis pipelines included: Python 3.12.1; NumPy 2.2.6, pandas 2.3.2, SciPy 1.16.3, statsmodels 0.14.6, and scikit-learn 1.7.2 for statistical analyses and predictive modeling; nibabel 5.4.0 for neuroimaging file input/output and neuromaps 0.0.7 for cortical-gradient and spin-test analyses; and Matplotlib 3.10.8 and Seaborn 0.13.2 for statistical visualization and nilearn 0.13.1 for fsLR-32k cortical-surface rendering, with Connectome Workbench 1.3.1 for cortical-surface rendering and border generation. Preprocessing used the ABCD-BIDS pipeline (https://github.com/DCAN-Labs/abcd-hcp-pipeline) and XCP-D version 0.13.0 (https://xcp-d.readthedocs.io/).

## Acknowledgements

We thank the participating families and ABCD research staff for their time and effort.

## Author Contributions Statement

Conceptualization (CA, IHG); Data curation (CA, EG, SK); Methodology (CA, EG, AZ, BJ, KMP, RAP); Formal analysis (CA, EG); Validation (CA, RAP); Writing – original draft (CA, EG, IHG); Writing – review and editing (CA, EG, GG, SK, JPU, JAR, YL, AZ, BJ, KMP, RAP, IHG); Funding acquisition (CA, EG, JPU, IHG).

