## Supplementary Material for "Early Adversity Selectively Reshapes the Somato-Cognitive Action Network in the Developing Brain"

**Contents**

**Supplementary Methods**

- Model specification, nesting, and convergence

**Supplementary Results**

- Representativeness of the analytic sample
- Individual dimensions of adversity and Bayes factors
- Associations with psychopathology (CBCL)

**Supplementary Tables**

- Table S1. Robustness of the threat–SCAN association across seven variance structures
- Table S2. Design and nesting structure of the analytic sample
- Table S3. Comparison of included and excluded participants
- Table S4. Associations of the ten adversity factors with SCAN size, with Bayes factors
- Table S5. Joint model of all ten adversity factors predicting SCAN size
- Table S6. SCAN mediation of threat and year-6 CBCL psychopathology
- Table S7. Within-person SCAN change and CBCL psychopathology

**Supplementary Methods**

Adversity exposure factors and composites

Early-life adversity was characterized by ten data-driven factors derived from the early-environment measures collected in the ABCD Study^1^, each z-scored across the analytic sample. Following the Dimensional Model of Adversity and Psychopathology^2,3^, we grouped the ten factors *a priori* into three composites, each computed as the mean of its constituent z-scored factors: threat (experiences of harm or its anticipation) – physical trauma, family aggression, family conflict, and family anger; deprivation (the absence of expected cognitive and social input) – socioeconomic and neighborhood disadvantage, low primary caregiver support, low secondary caregiver support, and caregiver supervision; and unpredictability (instability of the caregiving environment) – caregiver psychopathology and caregiver substance use or separation.

Three factors – family anger and primary and secondary caregiver support – were reverse-coded before compositing so that higher scores on every factor and every composite denote greater adversity. Each factor was positively associated with the composite to which it was assigned (*r* = 0.41-0.92; Figure S1). The three composites were themselves moderately intercorrelated (threat-deprivation *r* = 0.75; threat-unpredictability *r* = 0.73; deprivation-unpredictability *r* = 0.60), which motivates the multivariate model in which the composites compete to predict SCAN size (main text); Table S5 reports the corresponding model that enters all ten individual factors simultaneously.

Model specification, nesting, and convergence

All associations between early life adversity and cortical network topography, encroachment, and functional connectivity reported in the main text were estimated with ordinary least squares (OLS), adjusting for age, sex, and in-scanner head motion (mean framewise displacement), with study site entered as a set of fixed-effect indicators. Non-independence of siblings was accommodated with family-cluster-robust (sandwich) standard errors rather than a family random intercept.

We adopted this specification for three reasons, each documented in Table S2. First, all three scanner manufacturers and all 29 scanner serial numbers were perfectly nested within study site (no site used more than one manufacturer, and no serial number spanned more than one site), so fixed site indicators fully absorb between-scanner as well as between-site variance; adding a scanner random effect on top of site is therefore unidentified. Second, families never crossed sites (0 of 4,058), so family is structurally nested within site. Third, only 456 families (11.2%) contributed more than one imaged sibling, so a family random-intercept variance is estimated at essentially zero (its covariance is singular in every model that includes it). We therefore handle sibling non-independence with family-clustered robust standard errors rather than a boundary-valued variance component.

To confirm that this fixed-effects specification does not drive the findings, we refit the primary threat-SCAN association – both the single-predictor (bivariate) and the joint three-composite (multivariate) models – under seven variance structures (Table S1): (i) OLS with fixed site and family-cluster-robust standard errors (the specification reported throughout); (ii) OLS with fixed site and classical standard errors; (iii) a random site intercept; (iv) fixed site with a random family intercept; (v) crossed random site and family intercepts; (vi) a random site with family nested within site; and (vii) fixed site with an added scanner-serial random effect and a random family intercept. The standardized association was invariant across all seven specifications (bivariate partial *r* = 0.151-0.161; multivariate *β* = 0.00231-0.00233), and every model reached optimizer convergence. Specifications that include a family random intercept return a singular random-effects covariance, reflecting the near-zero family variance described above; this is a property of the sibling structure rather than a failure of estimation, and the fixed-effect inference on which all reported associations rest is unaffected.

**Supplementary Results**

Representativeness of the analytic sample

Relative to the remainder of the ABCD baseline cohort (n = 7,345), the analytic sample (n = 4,525) was modestly more likely to be female (52.0% vs. 45.3%), White (61.9% vs. 48.2%), and from higher-income and higher-education households (Table S3). All associations were small (bias-corrected^4^ Cramér's V = 0.065-0.164). This selection pattern is consistent with the well-documented association between usable MRI quality and sociodemographic characteristics in developmental neuroimaging^5^, and we note it as a bound on generalizability in the main-text limitations.

Individual dimensions of adversity and Bayes factors

The main text reports associations between the three adversity composites and SCAN size. Table S4 provides the parallel results for each of the ten individual adversity factors, together with Bayes factors. Nine of the ten factors were independently associated with SCAN size at baseline (all FDR *q* < .05); the exception was caregiver supervision. Bayes factors, computed as BIC approximations from the same fixed-site models (BF_10_ = exp[(BIC_null − BIC_full)/2]), corroborated this pattern: three of the four threat-type factors (physical trauma, family aggression, and family conflict) provided very strong evidence for an association (BF_10_ ranging from 5.7×10^10^ to 5.3×10^16^); family anger provided weaker but still positive evidence for an association (BF_10_ = 6.1); and only caregiver supervision, the one factor that was non-significant by FDR, favored the null (BF_01_ = 3.0).

Because the ten factors are correlated, we also fit a single model entering all ten simultaneously (Table S5). In this joint model, only family aggression remained independently associated with SCAN size after FDR correction (*β* = 0.0012, *q* < .001), consistent with substantial shared variance among the threat-type factors and supporting our decision to summarize them with the *a priori* threat, deprivation, and unpredictability composites rather than to interpret each factor in isolation.

**Associations with psychopathology (CBCL)**

We tested whether baseline SCAN size mediated the association between threat and each of 14 Child Behavior Checklist (CBCL) syndrome and DSM-oriented subscales at year 6, using the same bootstrapped (5,000 family-clustered resamples), covariate-matched mediation framework applied to cognition (Table S6). No subscale showed a significant indirect effect: none survived FDR correction and every bootstrap confidence interval included zero (all bootstrap *p* > .05; all FDR *q* > .80). Consistent with the cross-sectional models, within-person change in SCAN size did not track within-person change in any of the 14 CBCL subscales (none survived FDR correction; all *q* > .06; Table S7). Across both cross-sectional and within-person tests we found no evidence that SCAN size relates to concurrent or developing psychopathology, underscoring the specificity of the SCAN-cognition association reported in the main text.

**Supplementary Tables**

***Table S1.*** *Robustness of the threat-SCAN association across seven variance structures*

| Model | Specification | Estimate | *SE* | *p* | Converged |
| --- | --- | --- | --- | --- | --- |
| Bivariate | OLS, fixed site + family cluster-robust SE | partial *r* = 0.151 | 0.00021 | 1.1 × 10^-24^ | Yes |
| Bivariate | OLS, fixed site, classical SE | partial *r* = 0.158 | 0.00020 | 1.7 × 10^-26^ | Yes |
| Bivariate | MixedLM, fixed site + random family intercept | partial *r* = 0.158 | 0.00020 | 9.7 × 10^-27^ | Yes |
| Bivariate | MixedLM, random site intercept | partial *r* = 0.158 | 0.00019 | 7.1 × 10^-27^ | Yes |
| Bivariate | MixedLM, crossed random site + random family | partial *r* = 0.161 | 0.00017 | 5.2 × 10^-28^ | Yes |
| Bivariate | MixedLM, random site + family nested in site | partial *r* = 0.161 | 0.00017 | 5.2 × 10^-28^ | Yes |
| Bivariate | MixedLM, fixed site + random scanner-serial + random family | partial *r* = 0.158 | 0.00020 | 9.2 × 10^-27^ | Yes |
| Multivariate | OLS, fixed site + family cluster-robust SE | *β* = 0.00233 | 0.00032 | 2.8 × 10^-13^ | Yes |
| Multivariate | OLS, fixed site, classical SE | *β* = 0.00233 | 0.00030 | 1.3 × 10^-14^ | Yes |
| Multivariate | MixedLM, fixed site + random family intercept | *β* = 0.00231 | 0.00030 | 3.6 × 10^-14^ | Yes |
| Multivariate | MixedLM, random site intercept | *β* = 0.00233 | 0.00030 | 7.9 × 10^-15^ | Yes |
| Multivariate | MixedLM, crossed random site + random family | *β* = 0.00232 | 0.00030 | 8.8 × 10^-15^ | Yes |
| Multivariate | MixedLM, random site + family nested in site | *β* = 0.00232 | 0.00030 | 8.8 × 10^-15^ | Yes |
| Multivariate | MixedLM, fixed site + random scanner-serial + random family | *β* = 0.00232 | 0.00030 | 3.4 × 10^-14^ | Yes |

**Note.** Bivariate = single-predictor threat-SCAN model; Multivariate = threat estimate from the joint three-composite model. Specification (i), OLS with fixed site and family-cluster-robust standard errors, is the model reported throughout the main text. The standardized association is invariant across all seven specifications and every model converged.

***Table S2.*** *Design and nesting structure of the analytic sample*

| Quantity | Value | Interpretation |
| --- | --- | --- |
| Study-site indicator levels | 22 | sites entered as fixed-effect dummies |
| Scanner manufacturers | 3 | too few levels for a stable random-effect variance (<5-8) |
| Sites using >1 manufacturer | 0 | if 0, manufacturer is perfectly nested in site -> absorbed by site dummies |
| Scanner serial numbers | 29 | scanner serials |
| Serial numbers spanning >1 site | 0 | if 0, serial nested in site -> absorbed by site dummies |
| Maximum serial numbers per site | 2 | sites with multiple scanners over the study |
| Distinct families | 4058 | distinct family IDs in the sample |
| Families spanning >1 site | 0 | if 0, families never cross sites -> family is structurally nested in site |
| Families with ≥2 imaged siblings | 456 | 11.2% of families |
| Participants with a sibling in sample (%) | 20.4 | limited within-family replication -> family variance poorly identified |
| Models with singular family variance | 15/15 | singular family RE in the crossed models (family var at boundary) |

**Note.** The study-site variable comprises 22 indicator levels – the 21 ABCD research sites plus one administrative level (n = 15) for participants not affiliated with a standard collection site; all entered as fixed-effect indicators. Because scanner manufacturer and serial number are perfectly nested within site, and families never cross sites, fixed site indicators absorb the scanner and between-site variance; the near-zero family variance motivates family-cluster-robust standard errors in place of a family random intercept.

***Table S3.*** *Comparison of included and excluded participants*

| Characteristic | Excluded % | Included % | *Χ^2^* (df) | *p* | Cramér's *V* |
| --- | --- | --- | --- | --- | --- |
| Sex (% female) | 45.3 | 52.0 | χ^2^(1) = 50.9 | 9.6 × 10^-13^ | 0.065 |
| Race/ethnicity (% White) | 48.2 | 61.9 | χ^2^(4) = 258.2 | 1.1 × 10^-54^ | 0.153 |
| Household income (% ≥ $100k) | 41.5 | 54.9 | χ^2^(9) = 294.3 | 4.3 × 10^-58^ | 0.164 |
| Parental education (% ≥ bachelor’s) | 50.2 | 62.4 | χ^2^(20) = 211.6 | 5.5 × 10^-34^ | 0.133 |

**Note.** Excluded = remainder of the ABCD baseline cohort (n = 7,345); Included = analytic sample (n = 4,525). Percentages are computed on complete cases within each characteristic (participants missing that characteristic are omitted from its comparison) and therefore differ slightly from the corresponding Table 1 values, which are proportions of the full analytic sample (n = 4,525) with missing data shown separately. The χ^2^ tests and bias-corrected Cramér's V^4^ evaluate the full multi-category distribution of each characteristic – sex (2 levels), race/ethnicity (5), household income (10 categories), and parental education (21) – whereas the percentage columns report a single representative level: income ≥ $100k combines categories 9-10 and parental education ≥ bachelor's combines categories 18-21. All effect sizes are small.

***Table S4.*** *Associations of the ten adversity factors with SCAN size, with Bayes factors*

| Adversity factor | Partial *r* | FDR *q* | Bayes factor | Evidence |
| --- | --- | --- | --- | --- |
| Physical trauma | +0.131 | < .001 | BF_10_ = 5.3 × 10¹⁶ | very strong for H1 |
| Caregiver substance use / separation | +0.113 | < .001 | BF_10_ = 4.6 × 10¹¹ | very strong for H1 |
| Family conflict | +0.110 | < .001 | BF_10_ = 5.7 × 10¹⁰ | very strong for H1 |
| Family aggression | +0.106 | < .001 | BF_10_ = 1.8 × 10¹¹ | very strong for H1 |
| SES / neighborhood | +0.072 | < .001 | BF_10_ = 3.8 × 10³ | very strong for H1 |
| Primary caregiver support | -0.062 | < .001 | BF_10_ = 367 | very strong for H1 |
| Secondary caregiver support | -0.057 | .003 | BF_10_ = 46 | strong for H1 |
| Family anger | +0.051 | .011 | BF_10_ = 6.1 | positive for H1 |
| Caregiver psychopathology | +0.044 | .034 | BF_10_ = 1.9 | small for H1 |
| Caregiver supervision | +0.036 | .105 | BF_01_ = 3.0 | positive for H0 |

**Note.** Partial correlations of each adversity factor with baseline SCAN cortical share, adjusting for age, sex, head motion, and fixed site. FDR q corrected across the full 10-factor × 15-network matrix. Bayes factors are BIC approximations from the same models; BF_10_ > 1 favors an association, BF_01_ > 1 favors the null. Evidence labels follow Kass & Raftery (1995)^6^.

***Table S5.*** *Joint model of all ten adversity factors predicting SCAN size*

| Adversity factor | *β* | *SE* | *p* | FDR *q* |
| --- | --- | --- | --- | --- |
| Family Aggression | +0.0012 | 0.0002 | 5.1 × 10^-7^ | < .001 |
| Physical Trauma | +0.0011 | 0.0004 | .006 | .167 |
| SES/Neighborhood | -0.0013 | 0.0005 | .012 | .177 |
| Family Conflict | +0.0005 | 0.0003 | .049 | .326 |
| CG Psych | -0.0003 | 0.0002 | .108 | .447 |
| Family Anger | -0.0006 | 0.0004 | .110 | .447 |
| CG Substance/Sep | +0.0006 | 0.0004 | .188 | .542 |
| Primary CG Support | -0.0002 | 0.0002 | .329 | .665 |
| CG Supervision | +0.0001 | 0.0002 | .627 | .856 |
| Secondary CG Support | -0.0001 | 0.0002 | .722 | .914 |

**Note.** All ten adversity factors entered simultaneously into a single OLS model predicting baseline SCAN cortical share (covariates and fixed site as above). Only family aggression remains independently associated after FDR correction, reflecting shared variance among the threat-type factors and supporting the composite approach.

***Table S6.*** *SCAN mediation of threat and year-6 CBCL psychopathology*

| CBCL subscale | Indirect effect | 95% CI | Bootstrap *p* | FDR *q* |
| --- | --- | --- | --- | --- |
| Total Problems | +0.009 | [-0.099, 0.130] | .897 | .953 |
| Internalizing | +0.018 | [-0.026, 0.068] | .458 | .953 |
| Externalizing | -0.004 | [-0.042, 0.033] | .797 | .953 |
| Anxious/Depressed | -0.002 | [-0.023, 0.019] | .811 | .953 |
| Withdrawn/Depressed | +0.003 | [-0.015, 0.023] | .768 | .953 |
| Somatic Complaints | +0.013 | [-0.004, 0.034] | .132 | .927 |
| DSM-5 Depressive | -0.008 | [-0.031, 0.015] | .447 | .953 |
| Aggressive Behavior | -0.007 | [-0.030, 0.014] | .512 | .953 |
| Rule-Breaking | +0.001 | [-0.018, 0.020] | .923 | .953 |
| DSM-5 Conduct | -0.007 | [-0.024, 0.009] | .390 | .953 |
| Attention Problems | +0.005 | [-0.017, 0.029] | .669 | .953 |
| DSM-5 ADHD | -0.000 | [-0.017, 0.017] | .953 | .953 |
| Thought Problems | -0.014 | [-0.030, 0.000] | .059 | .829 |
| Social Problems | -0.005 | [-0.016, 0.006] | .418 | .953 |

**Note.** Indirect effect of threat on each year-6 CBCL subscale through baseline SCAN size (n = 1,513), bootstrapped over 5,000 family-clustered resamples with a matched baseline-subscale covariate. Negative indirect effects indicate that a larger SCAN is associated with fewer problems.

***Table S7.*** *Within-person SCAN change and CBCL psychopathology*

| CBCL subscale | Within-person *β* | *p* | FDR *q* |
| --- | --- | --- | --- |
| Total Problems | -0.009 | .526 | .761 |
| Internalizing | -0.029 | .048 | .212 |
| Externalizing | +0.012 | .369 | .738 |
| Anxious/Depressed | -0.025 | .076 | .213 |
| Withdrawn/Depressed | -0.041 | .005 | .064 |
| Somatic Complaints | -0.004 | .768 | .868 |
| DSM-5 Depressive | -0.031 | .041 | .212 |
| Aggressive Behavior | +0.008 | .544 | .761 |
| Rule-Breaking | +0.016 | .283 | .660 |
| DSM-5 Conduct | +0.026 | .061 | .212 |
| Attention Problems | +0.006 | .667 | .848 |
| DSM-5 ADHD | -0.000 | .995 | .995 |
| Thought Problems | -0.009 | .531 | .761 |
| Social Problems | -0.003 | .806 | .868 |

**Note.** All-waves within-person models (n = 6,094 observations from 3,442 children) relating within-person change in SCAN size to within-person change in each of the 14 CBCL subscales. No subscale tracked within-person SCAN change (none survived FDR correction; all *q*s > .06

**Supplementary Figures**

***Figure S1.*** *Intercorrelations among the ten adversity factors and the three composites*

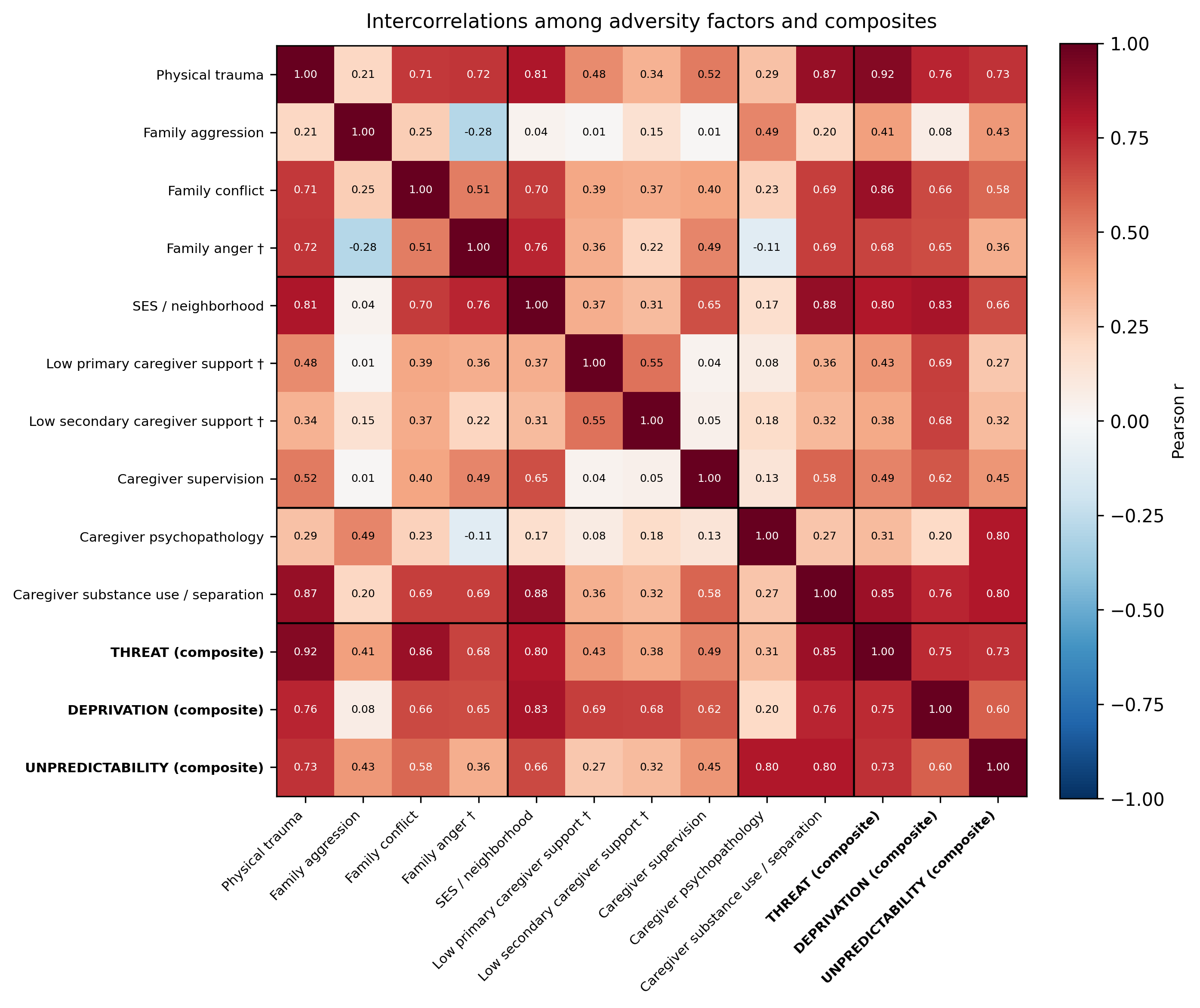
 **Note***.* Pearson correlations among the ten z-scored adversity factors (grouped by composite) and the threat, deprivation, and unpredictability composites, in the analytic sample (n = 4,525). Factors are shown in composite-aligned orientation, so that higher values on every factor denote greater adversity; † marks the three factors reverse-coded to achieve this alignment (family anger, primary and secondary caregiver support). Black lines delineate the composite groupings. Each factor correlates positively with the composite to which it was assigned (*r* = 0.41-0.92), and the composites are themselves moderately intercorrelated, motivating the multivariate specification used in the main text.
